# Methylation-aware long-read phasing significantly improves genome-wide haplotype reconstruction

**DOI:** 10.64898/2026.03.11.710820

**Authors:** Aaron Pfennig, Joshua M. Akey

## Abstract

Haplotypes are linear sequences of co-inherited alleles along individual chromosomes and are central to genetic mapping, clinical variant interpretation, and inference of population history. However, accurate genome-wide haplotype reconstruction remains challenging. Long-read sequencing has the potential to dramatically improve haplotype inference, but existing methods do not directly leverage all the information embedded in these data. Here, we present LongHap, a read-based phasing method that integrates sequence and 5-methylcytosine (5mC) information in a unified probabilistic framework. By leveraging differentially methylated sites, LongHap resolves phase relationships between variants that are inaccessible to sequence-based approaches alone. Across multiple datasets and sequencing platforms, LongHap increases phase block lengths by up to 30% while substantially reducing switch error rates. LongHap rigorously embeds complex structural variants into the broader haplotype context using loopy belief propagation, enabling improved phasing of INDELs and other variant classes that are inherently difficult to resolve. Methylation-aware phasing also improves the accuracy and contiguity of haplotypes spanning rare variants and structurally complex, medically relevant genes across diverse ancestries, facilitating the interpretation of compound heterozygosity and haplotype-specific regulatory architectures. These results establish methylation-aware phasing as a general framework for improving genome-wide haplotype reconstruction, with broad applications across genetics and genomics.

## Introduction

Genetic variation is inherently organized into haplotypes—combinations of alleles that are inherited together on individual chromosomes—and many fundamental biological processes operate at this level rather than at individual variant sites^1^. Haplotype structure shapes patterns of linkage disequilibrium, governs the joint effects of multiple variants on gene regulation and protein function, and is critical for diverse applications ranging from association mapping and clinical variant interpretation to inference of population history and natural selection^2,3^.

To facilitate powerful haplotype-based interpretation of genetic variation, phasing approaches have been developed that fall into three main categories: statistical, pedigree (trio), and read-based phasing^4^. Using large reference panels and parental sequencing data, statistical and trio phasing can infer haplotypes on a chromosome-scale^5,6^, but rare variant phasing remains challenging, particularly in complex genomic regions, since statistical inference is underpowered and short reads are difficult to map in these regions^7,8^. Read-based phasing, in contrast, directly analyzes raw sequencing data to identify alleles that co-occur across heterozygous sites within individual reads^9–13^, avoiding reference panel biases and enabling rare variant phasing without requiring parental sequencing data^7^. However, the ability of read-based approaches to phase variants over long distances has historically been limited by short read lengths.

Improved accuracy and declining costs have made long-read sequencing with PacBio and Oxford Nanopore Technologies (ONT) viable alternatives to short-read sequencing ^14,15^. The longer read lengths of these technologies dramatically improve the ability of read-based phasing approaches, such as WhatsHap, HapCUT2, and HaploMaker, to infer accurate haplotypes over longer genomic segments^9–12^. A recent method, called LongPhase, achieves chromosome-scale phasing with ultra-long ONT (UL-ONT) data, while improving phasing contiguity and computational efficiency over WhatsHap and HapCUT2^13^. However, LongPhase relies on simplified modeling of read evidence, which can limit its ability to robustly phase complex variants and regions affected by alignment uncertainty. In addition to DNA sequence variation, modern long-read sequencing technologies capture additional layers of information, including base modifications such as 5-methylcytosine (5mC), which can provide orthogonal evidence of haplotype identity. Despite the availability of epigenetic information in long-read sequencing data, only one method, MethPhaser^16^, has been developed to harness 5mC data in long reads to *post hoc* refine the haplotypes inferred by another read-based phasing tool. Thus, no read-based phasing method exists that integrates sequence and epigenetic information in a unified framework, leaving orthogonal signals in long-read sequencing unexploited.

To address these limitations, we developed LongHap, a read-based phasing method that seamlessly integrates sequence and 5mC methylation information from PacBio HiFi and ONT sequencing data in a unified framework (Figure 1). By integrating methylation signals, LongHap resolves phase relationships that cannot be inferred from sequence data alone. This enables the extension of phase blocks across gaps lacking read overlap, reducing switch errors by up to 5% and increasing phase block contiguity by 30%. Crucially, LongHap’s novel embedding of structural variants (SVs) into the broader haplotype context through loopy belief propagation extends the benefits of methylation-aware phasing to all variant classes. In addition to these genome-wide performance gains, we demonstrate that integrating methylation information enables phasing of challenging, medically relevant genes and rare variants that are difficult to phase otherwise, with direct implications for interpreting compound heterozygosity in clinical settings^5,17,18^. These results highlight the power of exploiting the full information content of long-read sequencing data rather than simply extending short-read paradigms to long-read data. The substantial improvements in phasing accuracy and contiguity afforded by LongHap will facilitate inferences into the biology and evolution of complex eukaryotic genomes.

**Figure 1.**
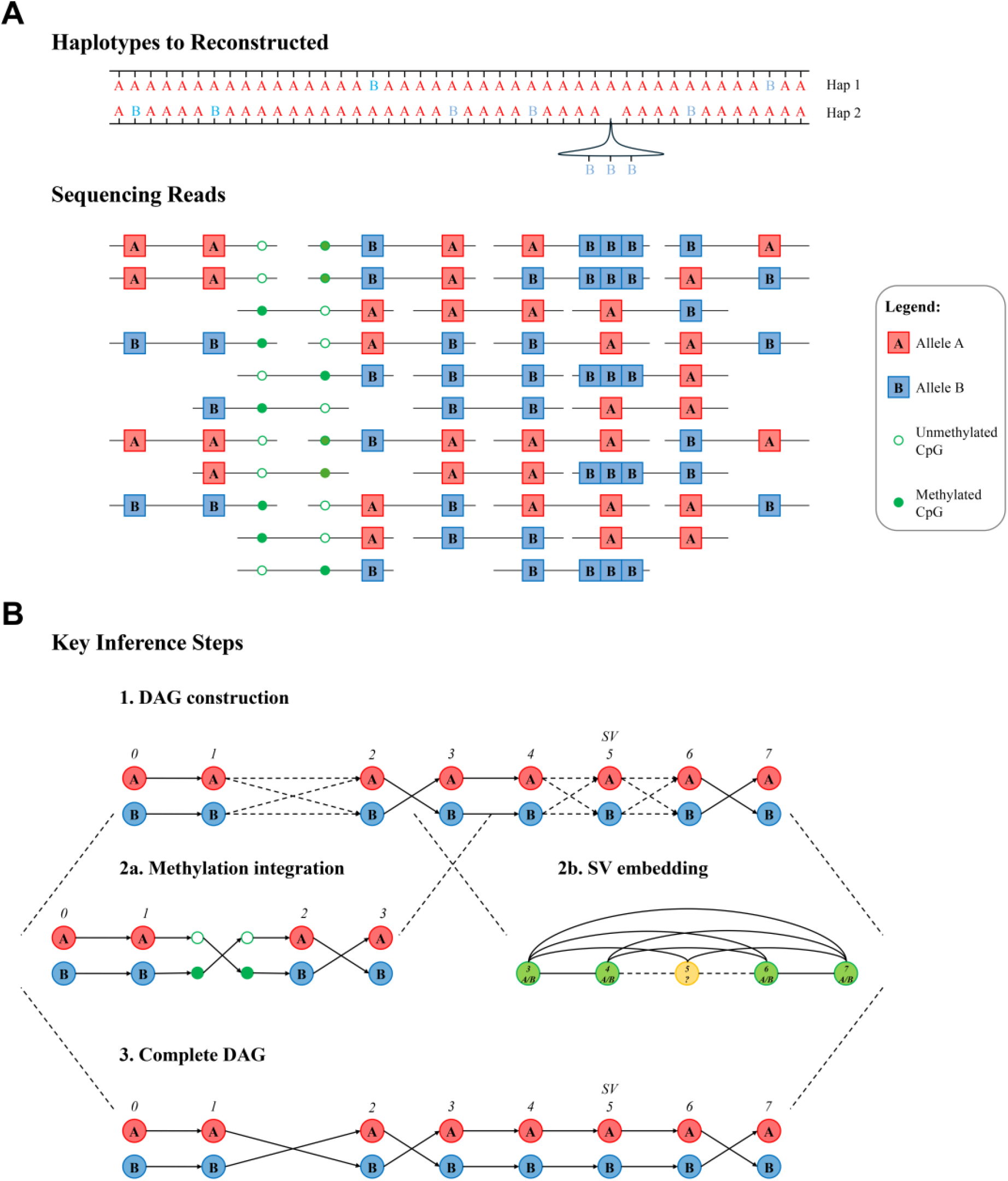
Schematic overview of LongHap. **A)** The haplotype structure at a locus that we want to reconstruct from long-read sequencing data. A and B denote alleles, and the bubble represents a three-base pair insertion on haplotype 2. Open and filled green circles indicate unmethylated and methylated CpG sites, respectively. Sequencing reads across the region are shown below the haplotypes. **B)** Schematic of the three key steps LongHap uses to infer haplotypes from read-based sequence and methylation variation. LongHap constructs a directed acyclic graph (DAG) in which each layer represents a heterozygous variant and edge weights reflect the co-occurrence of alleles across individual sequencing reads. Where adjacent phase blocks cannot be connected by sequence evidence alone, LongHap iteratively identifies differentially methylated sites on-the-fly and uses them to assign reads to maternal and paternal haplotypes, resolving transitions between two previously ambiguous layers. Finally, for complex or low-support variants, such as the structural variant at position 5 in the example shown, LongHap extracts a local subgraph and applies loopy belief propagation to incorporate long-range haplotype context, re-estimating transition probabilities into and out of the target variant. The complete DAG is then decoded using a Viterbi-like algorithm to identify the most likely haplotype configuration.

## Results

### Overview of LongHap

LongHap is a read-based variant phasing method that integrates methylation signals native to long-read sequencing data, such as PacBio Revio HiFi and ONT sequencing, into a unified framework. Conceptually, LongHap reframes haplotype phasing as a problem of joint probabilistic inference from multiple, partially independent signals embedded within sequencing reads. As input, LongHap requires variant calls in VCF format, aligned sequencing reads in BAM format, and unphased methylation calls (Figure 1A). LongHap then uses a stepwise approach to phase SNVs, INDELs, and SVs (Figure 1B). First, LongHap phases pairs of variants based on sequence information. In a second step, LongHap dynamically identifies differentially methylated sites and leverages them as additional, phase-informative markers to resolve variant pairs that could not be confidently phased based on sequence information alone, thereby extending initially inferred phase blocks. LongHap then embeds complex and low-support variants into the broader haplotype context using graph theory, that is, loopy belief propagation, for more robust phasing (Figure 1B). Finally, LongHap outputs a phased VCF and, optionally, haplotagged read alignments, phase block coordinates, and the set of differentially methylated sites used for phasing.

### LongHap outperforms all existing tools

We evaluated LongHap’s performance using PacBio Revio HiFi and regular and ultra-long ONT long-read sequencing datasets for HG002 that were made available by the T2T consortium and Human Pangenome Reference Consortium (HPRC)^19,20^. For all autosomes, we calculated the switch error rates, the percentage of phased heterozygous sites, and phase block length N50s for LongHap, WhatsHap, HapCUT2, LongPhase, and WhatsHap + MethPhaser using WhatsHap compare and stats^9–11,13,16^. To increase comparability and highlight the benefits of integrating methylation information, we compare LongHap’s performance to the other tools without and with methylation information. No tool phases all heterozygous sites, and we thus limit the comparisons to the intersection of biallelic sites that were phased by all tools.

A key challenge in read-based phasing is simultaneously minimizing switch errors, while maximizing the fraction of variants phased. LongHap advances this trade-off, outperforming existing tools on this combined criterion across long-read sequencing technologies. Using PacBio Revio HiFi data, LongHap achieves a switch error rate of 0.196% with methylation information, while phasing 99.13% of heterozygous sites (Figures 2A & B). In contrast, LongPhase achieves a marginally lower switch error rate (0.192%) but phases 18,394 fewer sites (98.42%). Conversely, WhatsHap and HapCUT2 phase more sites but with considerably higher error rates (~0.246%) (Figure 2B). Notably, LongHap’s integration of methylation information results in 164 fewer errors than sequence-only phasing. In contrast, MethPhaser, the only other tool that incorporates methylation, increases the switch error rate (0.281%) (Figure 2A & B), underscoring the advantage of jointly modeling sequence and epigenetic signals over *post hoc* refinement. As discussed below, the importance of balancing the switch error rate with the fraction of phased heterozygous sites is exacerbated for noisier ONT sequencing data.

**Figure 2.**
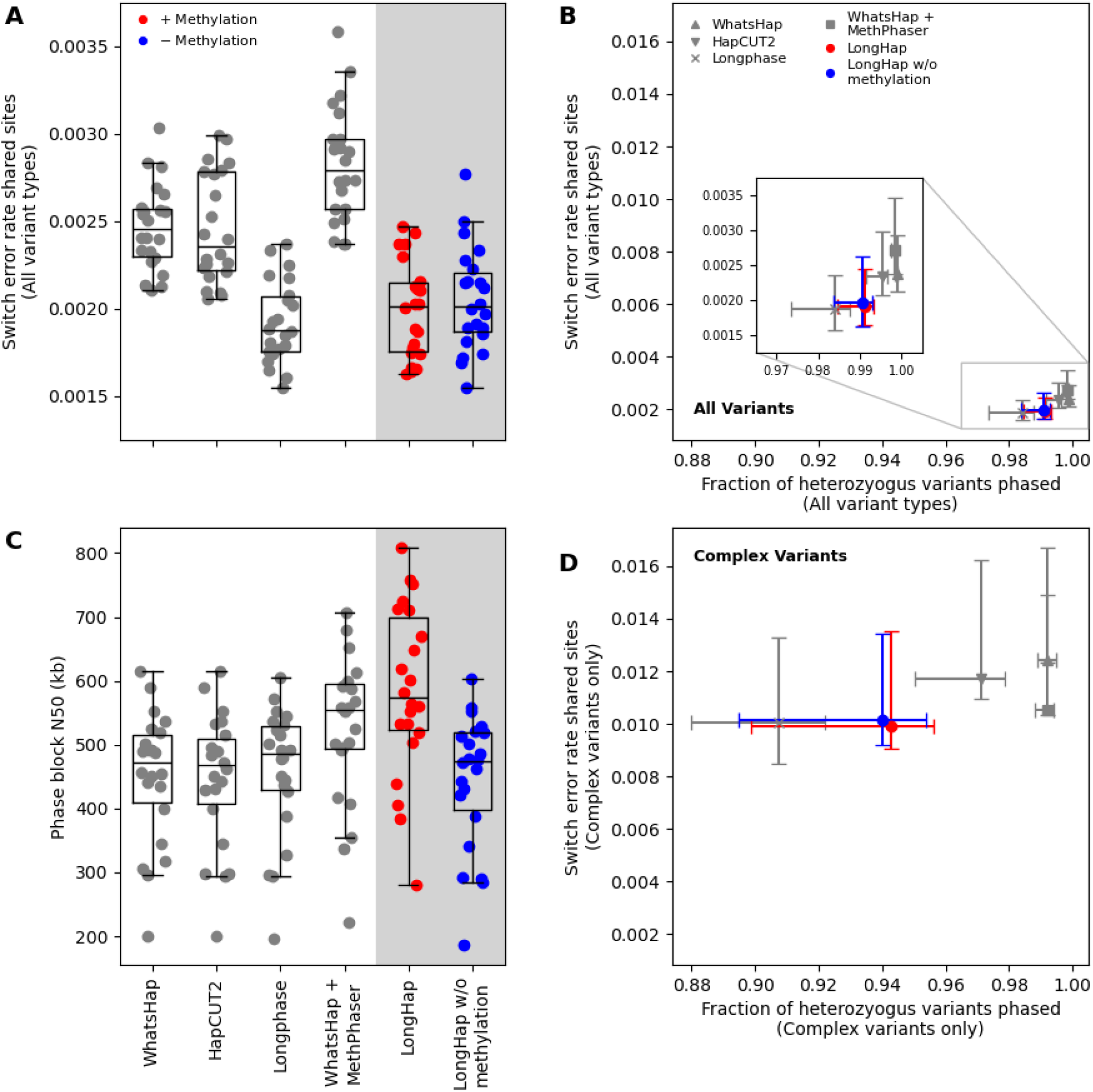
LongHap optimizes the trade-off between minimizing the switch error rate and maximizing the fraction of phased variants. **A)** Switch error rates for different read-based variant phasing tools using 38x PacBio Revio HiFi data for individual chromosomes of the well-characterized genome of HG002. LongHap makes 164 fewer errors when considering methylation information (4,860 vs. 5,024), and 2,117 fewer errors than WhatsHap + MethPhaser (6,977), another tool that integrates methylation information. LongPhase makes the fewest errors (4,754) but phases the fewest sites of all tools. Switch error rates were calculated based on the intersection of sites phased by all tools (96.13% of all heterozygous sites). **B)** By leveraging methylation information, LongHap phases more variants than LongPhase, while maintaining a lower error rate than all other tools. Switch error rates were calculated based on the intersection of heterozygous sites phased by all tools (96.13% of all heterozygous sites). **C)** Simultaneously, LongHap’s integration of methylation information significantly increases the phase block N50 (mean N50: 584 kb compared to 443 kb), a greater gain than achieved by WhatsHap + MethPhaser (mean N50: 523 kb compared to 450 kb for WhatsHap alone). **D)** Similar to **(B)** LongHap phases more INDELs and SVs than LongPhase, while maintaining a lower error rate than all other tools, through the integration of methylation information and its rigorous embedding of INDELs and SVs into the broader haplotype context. Switch error rates were calculated based on the intersection of INDEL and SV sites phased by all tools (84.80% of all heterozygous sites with complex variants).

Beyond switch error rates, phase block contiguity determines how much of the genome can be interpreted in a haplotype-resolved manner. While all methods achieve similar contiguity without methylation data (~443 kb), incorporating methylation signals increases LongHap’s phase block N50 by 31.9% to 584 kb, outperforming WhatsHap + MethPhaser (523 kb; +16.2%) (Figure 2C). In summary, these results demonstrate that integrating methylation information enables LongHap to extend phase blocks and resolve phase relationships that cannot be inferred from sequence data alone. This leads to a 3.3% reduction in genome-wide switch errors and a >30% increase in phase block contiguity.

### LongHap accurately co-phases complex structural variants

Complex variants such as INDELs and SVs are inherently more difficult to phase than SNVs due to sequencing errors and alignment ambiguities around breakpoints. LongHap addresses these challenges by co-phasing INDELs and SVs along with SNVs, combining local realignment to resolve alignment uncertainty with loopy belief propagation to efficiently embed complex variants into the broader haplotype context. In contrast, WhatsHap and HapCUT2 are primarily designed for SNV phasing^9–11^, while LongPhase treats INDELs and SVs similarly to SNVs, without explicitly modeling alignment uncertainty or complex variant structure^13^.

To assess the benefits of this principled approach, we separately evaluated the accuracy of each tool for INDELs and SVs. As with genome-wide benchmarks, LongHap advances the Pareto frontier for complex variant phasing. Without methylation information, LongHap achieves a switch error rate comparable to LongPhase (1.01% vs 1.00%), while phasing substantially more complex variants than LongPhase (94.29% vs 90.73%). Crucially, with methylation information, LongHap attains the lowest switch error rate of all tools (0.99%) and phases 12,929 (3.56%) more INDELs and SVs than LongPhase (Figure 2D). This demonstrates that methylation information is particularly valuable for complex variant phasing, where sequence evidence alone is most likely to be ambiguous or insufficient around variant breakpoints.

### LongHap capitalizes on longer read lengths of ONT sequencing data

Integrating methylation information yields consistent improvements across all ONT sequencing platforms, demonstrating that LongHap’s methylation-aware framework generalizes across modern long-read sequencing technologies (Figures S1 & S2). With ONT data, LongHap achieves a lower switch error rate than WhatsHap, HapCUT2, and WhatsHap + MethPhaser (0.110% without and with methylation information compared to 0.129%, 0.135%, and 0.141%, respectively) (Figure S1A), while phasing substantially more heterozygous sites than LongPhase (98.37% - 99.43% for LongHap compared to 93.81% - 96.42% for LongPhase) at a slightly higher switch error rate (0.110% vs. 0.081%) (Figure S1B). The difference between the tools becomes most acute for INDELs and SVs, with LongPhase phasing only 74.93% and 70.74% of heterozygous INDELs and SVs compared to 94.85% and 95.04% for LongHap with ONT and UL-ONT data, respectively (Figures S1D, S2D).

LongHap also successfully capitalizes on the greater read length in ONT and UL-ONT sequencing data, achieving chromosome-scale phasing (average phase block N50s: 1.97 Mb and 80.7 Mb, respectively). In contrast, WhatsHap fails to translate the longer read lengths into meaningfully longer phase blocks (average phase block N50s: 1.47 Mb and 5.82 Mb), and MethPhaser cannot compensate for WhatsHap’s poor phase block N50 (average phase block N50s: 1.69 Mb and 7.22 Mb) (Figures S1C & S2C).

Together, these data highlight the fundamental trade-off between accuracy and completeness in haplotype inference, whereby methods that minimize switch error rates often do so by excluding difficult-to-phase variants. LongHap optimizes this trade-off, phasing more variants than LongPhase but fewer than WhatsHap or HapCUT2, while maintaining an intermediate switch error rate with ONT and UL-ONT sequencing data.

### Gap-bridging properties of phase-informative methylation signals

Leveraging methylation information, LongHap connects 1,636 out of 9,005 phase blocks inferred based on sequence information (18.2%) that could not be resolved by sequence evidence alone using PacBio Revio HiFi data. The average gap between two connected blocks is 32 kb (median: 29 kb; maximum: 233 kb; standard deviation: 12 kb) (Figure 3A). On average, LongHap considers 20.2 differentially methylated sites (median: 15; SD: 19.9) to connect two phase blocks (Figure 3B). These gaps bridged between two phase blocks scale with read length. For ONT and UL-ONT sequencing data, the average distances between two phase blocks connected with differentially methylated sites are 56 kb (median: 51 kb; maximum: 325 kb; standard deviation: 24 kb) and 415 kb (median: 239 kb; maximum: 2,579 kb; standard deviation: 626 kb), respectively (Figures S3 & S4).

**Figure 3.**
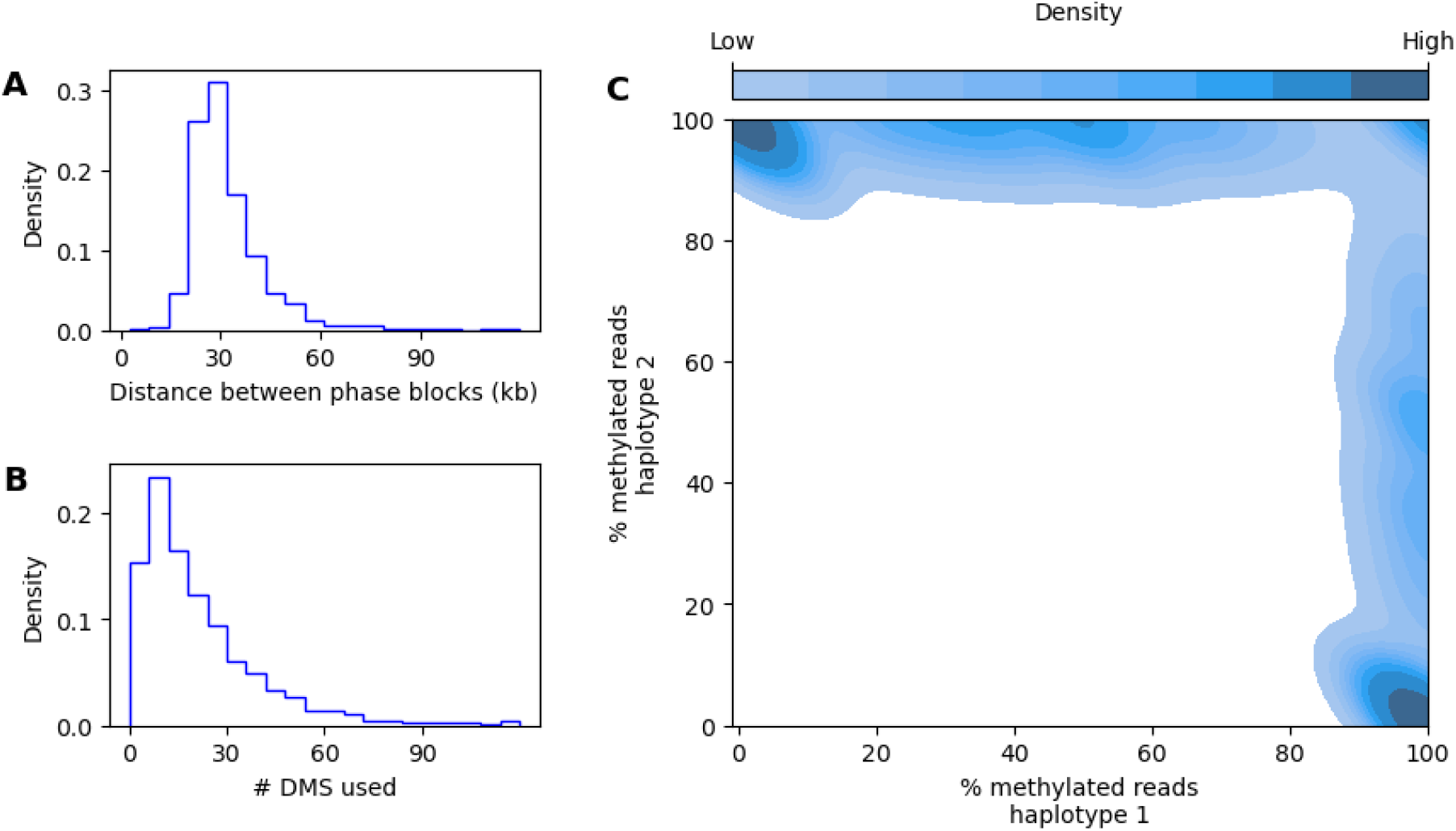
Characterization of differentially methylated sites (DMS) used by LongHap to bridge previously discontinuous phasing blocks. **A)** Distribution of distances between phase blocks connected using methylation information, using 38x PacBio Revio HiFi data. Most bridged gaps are between 20 and 40 kb, with a small number of gaps exceeding 120 kb. **B)** Distribution of the number of DMS sites used to bridge each gap. Most gaps are resolved using 20 DMS sites, with diminishing frequency at higher counts. **C)** Contour plot the fraction of reads with methylation marks assigned to either haplotype at sites used for phasing by LongHap. Most sites show clear signals of differential methylation, that is, they are truly phase-informative, as most reads are methylated on one haplotype while most reads are not methylated on the other haplotype (top left and bottom right corners). However, LongHap also identifies some false-positive differentially methylated sites (top right corner), injecting noise into the methylation-based phasing. A read is methylated at a given site if *P*(*meth*) > 0.5.

Phase-informative methylation sites are characterized by strongly asymmetric methylation patterns between haplotypes. To confirm that the differentially methylated sites leveraged by LongHap are informative for phasing, we compared the fraction of reads with evidence of methylation, *P*(*meth*) > 0.5, that support each haplotype. Most sites used by LongHap show differential methylation patterns with no or only an intermediate fraction of reads assigned to one haplotype being methylated, while most reads assigned to the other haplotype are unmethylated (Figure 3C).

### Methylation information enables the phasing of biomedically important loci

To evaluate LongHap’s ability to phase a recently curated list of 273 challenging, medically relevant genes (CMRGs)^21^, we intersected this gene list with phase block coordinates for each assessed tool. Integrating methylation information substantially improves LongHap’s phasing of CMRGs across all metrics. While LongHap and WhatsHap + MethPhaser both fully phase the most CMRGs (211), LongHap achieves the most contiguous phasing of the 247 CMRGs covered by all tools, with the fewest overlapping phase blocks (633 compared to 756 for HapCUT2) and the greatest coverage of phased CMRG base pairs (128.9 Mb compared to 127.3 Mb for WhatsHap) (Table 1). Strikingly, methylation integration alone reduces the number of phase blocks overlapping CMRGs from 714 to 633 (11.3%), representing a direct improvement in the interpretability of these clinically important regions. Without methylation information, LongHap’s CMRG phasing is comparable to or slightly worse than other tools, underscoring that methylation is the critical driver of these improvements. Similar patterns are observed with ONT and UL-ONT sequencing data (Tables S2 & S3).

**Table 1.** Number of phase blocks inferred by each tool using 38x PacBio Revio HiFi data that overlap challenging, medically relevant genes (CMRGs). 247 CMRGs are covered by all tools. LongHap infers the most contiguous phase blocks overlapping CMRGs, as indicated by the lowest number of phase blocks and the second most overlapping base pairs.

|  | CMRGs covered | Phase blocks overlapping CMRGs | CMRGs fully phased | Common phase blocks overlapping CMRGs | Overlapping base pairs of common CMRGs (Mb) |
| --- | --- | --- | --- | --- | --- |
| LongHap | 250 | 635 | 211 | 633 | 128.9 |
| LongHap w/o methylation information | 248 | 714 | 206 | 714 | 125.2 |
| LongPhase | 254 | 741 | 209 | 764 | 127.2 |
| WhatsHap | 254 | 774 | 210 | 781 | 127.3 |
| HapCUT2 | 253 | 759 | 209 | 756 | 126.6 |
| WhatsHap + MethPhaser | 254 | 873 | 211 | 883 | 126.7 |

To illustrate how adding methylation information improves phasing of CMRGs, we analyzed the *LIX1* gene in more detail (CHM13 coordinates: chr5:97,592,630-97,643,516; Figure 4). *LIX1* is expressed in the central nervous system and thyroid in humans and is involved in cell regulation and differentiation, as well as mitochondrial metabolism^22–25^. *LIX1* has recently been associated with worse prognoses for individuals with gastrointestinal stromal tumors^26^. Without methylation information, LongHap cannot phase the regions spanning the *LIX1* gene, as coverage dropout causes the phase block to break, leaving the gene split across two disconnected blocks. Integrating methylation information allows connecting the two adjacent phase blocks by leveraging phase-informative differentially methylated sites to assign additional reads to distinct haplotypes, producing a single continuous phase block spanning the entire locus (Figure 4).

**Figure 4.**
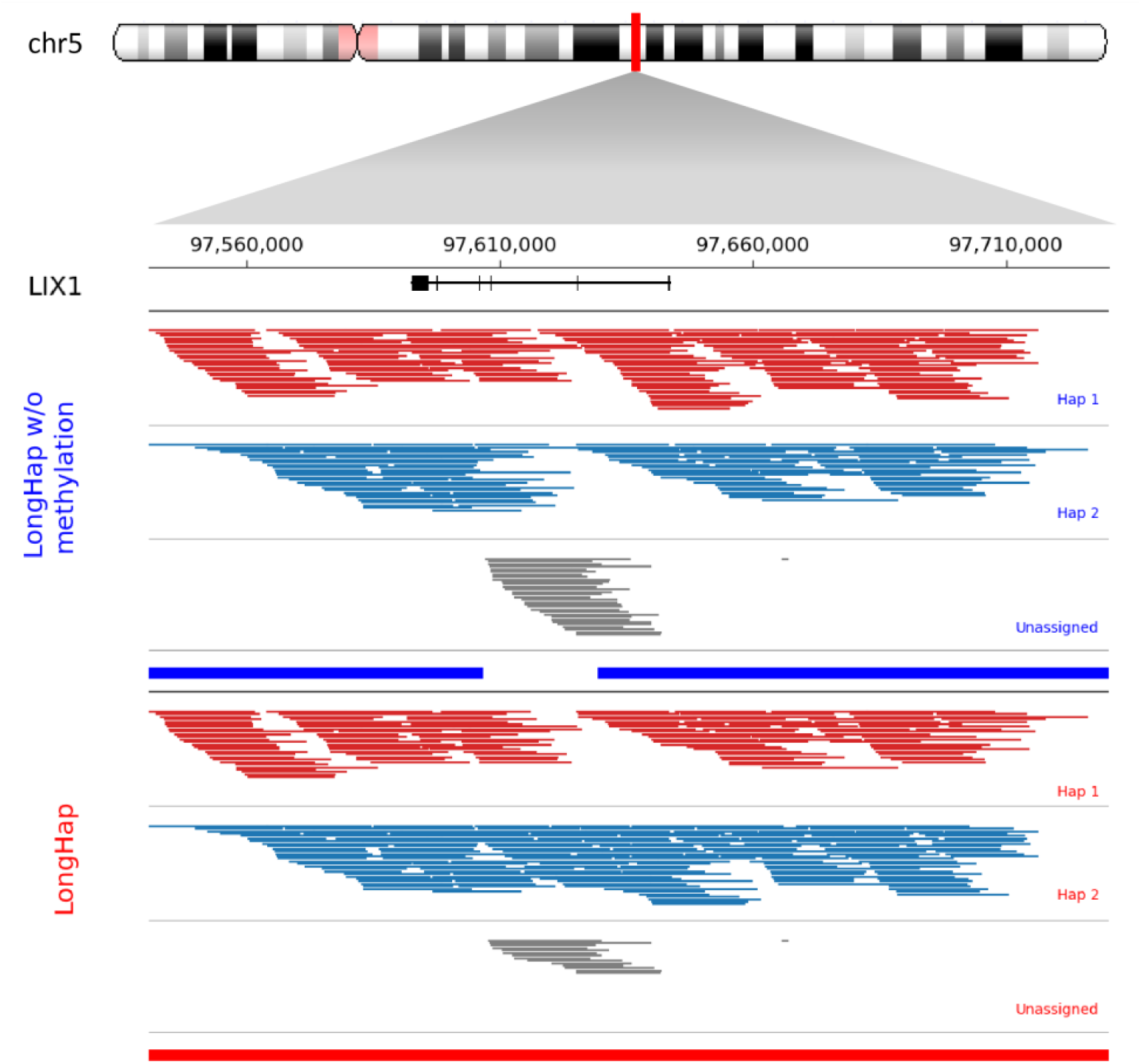
Methylation information enables continuous phasing across the LIX1 locus. Haplotagged read alignments at the LIX1 locus (chr5:97,592,630–97,643,516; CHM13v2.0) for LongHap run without (top) and with (middle) methylation information. Reads assigned to haplotype A are shown in red, haplotype B in blue, and unassigned reads in grey. Without methylation information, a coverage dropout causes the phase block to break into two. By leveraging differentially methylated sites to assign additional reads to distinct haplotypes, LongHap resolves a continuous phase block spanning the entire locus. Inferred phase blocks in the bottom tracks for LongHap without methylation (blue) and with methylation information (red). Genomic coordinates are shown relative to CHM13v2.0.

### Genome-wide inference of haplotypes in globally diverse individuals

We investigated LongHap’s ability to generate new biological knowledge by analyzing seven samples from diverse genetic ancestries using PacBio Revio HiFi data from the HPRC^20^. Specifically, we analyzed two Southern Han Chinese samples (HG00609, HG00658), two admixed American samples from Puerto Rico (HG00738, HG01099), and two sub-Saharan African Mandinka samples (HG02723, HG02615) in addition to HG002, which is of European descent. When comparing LongHap’s phasing against the *de novo* genome assemblies from HPRC release 2, LongHap achieves low switch error rates for all samples, ranging from 0.185% to 0.486% per chromosome. For the East Asian samples, the switch error rates are higher (0.313% - 0.486%) and inferred phase block N50s are shorter (mean: 321 kb) than for African samples (switch error rates: 0.199% - 0.374% per chromosome; mean phase block N50: 1,727 kb). This pattern reflects the higher heterozygosity of African samples, in which the greater density of heterozygous sites enables more contiguous phasing. However, the integration of methylation leads to the same relative improvements in phasing accuracy and contiguity independent of genetic ancestry (Figure S5).

Rare variants (MAF < 1%) are particularly challenging to phase using statistical approaches, which are underpowered at low allele frequencies, making read-based phasing essential for their accurate haplotype assignment^8^. By leveraging methylation signals, LongHap reduces switch errors between rare variants and increases the number of rare variants that can be phased. Specifically, using PacBio Revio HiFi data, LongHap makes, on average, 45 fewer errors when phasing rare variants per genome (range: −201 - +8; Figure 5A), while phasing substantially more rare variants (range 151 – 822) (Figure 5B). This underscores that the benefit of methylation-aware phasing is the largest in individuals of European and East Asian ancestry, where lower heterozygosity creates more frequent gaps in sequence-based phasing that methylation information can bridge.

**Figure 5.**
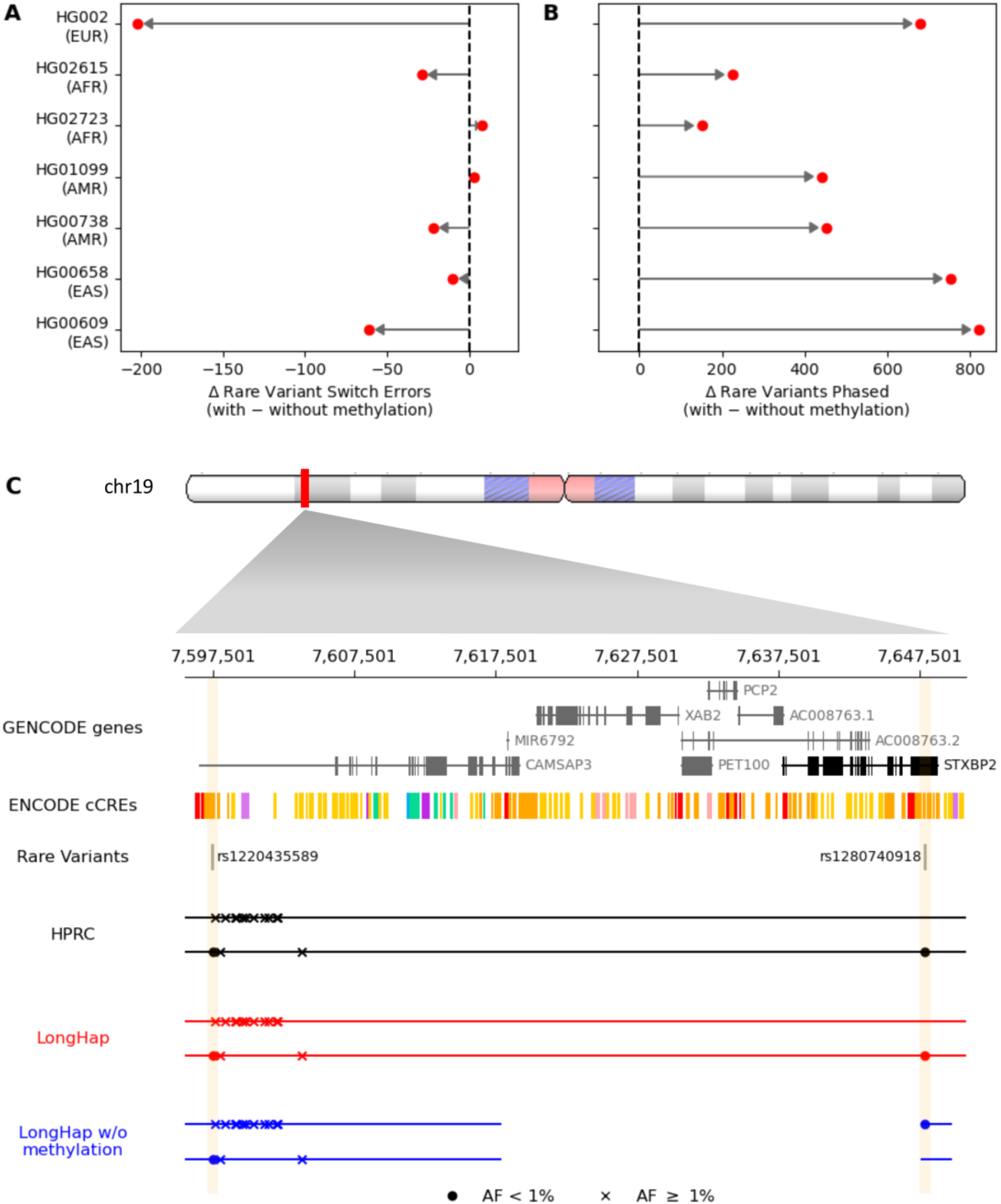
Methylation information improves phasing of rare variants across diverse ancestries and enables haplotype inference at a disease-relevant locus. **A)** Change in the number of switch errors between rare variants (MAF < 1%) when integrating methylation information across seven individuals of diverse genetic ancestries. Negative values indicate fewer errors with methylation. **B)** Corresponding change in the number of rare variants successfully phased. In all individuals, methylation integration reduces switch errors and increases the number of phased rare variants. **C)** At the STXBP2 locus in HG00609, LongHap without methylation information (bottom) places two rare variants (rs1220435589, rs1280740918) in separate phase blocks, preventing inference of their cis/trans configuration. Integrating methylation information (middle) resolves a continuous phase block, correctly identifying that both variants occur in cis in enhancer elements linked to STXBP2 expression. The HPRC assembly-based phasing (top) is shown as ground truth. Only variants present in the 1000 Genomes Project reference panel are shown. Genomic coordinates are shown relative to CHM13v2.0.

To illustrate this, we examined the *STXBP2* locus in HG00609, a CMRG implicated in Hemophagocytic Lymphohistiocytosis type 5, a fatal autoimmune disease (CHM13 coordinates: chr19: 7,637,764-7,648,756; Figure 5C)^27–31^. Without methylation information, two rare variants at this locus (rs1220435589, rs1280740918) fall in separate phase blocks, making it impossible to determine their *cis*/*trans* configuration. Consistent with the HPRC assembly, LongHap resolves a continuous phase block spanning both variants and infers that these variants occur in *cis* when leveraging methylation information, which is supported by the HPRC assembly, and is critical for interpreting their combined functional impact, as both variants reside in enhancer elements linked to *STXBP2* expression^32^. This example illustrates how LongHap can resolve clinically critical haplotypes, such as the haplotype-specific regulatory architecture at the STXBP2 locus, that are inaccessible to sequence-only approaches.

## Discussion

LongHap extends read-based phasing by leveraging methylation signals to resolve haplotype relationships that are inaccessible to sequence information alone, enabling more accurate and contiguous phasing, particularly for rare and structurally complex variants. Integrating methylation reduces switch error rates by up to 5% and increases phase block N50 by over 30%, translating into hundreds fewer phasing errors and thousands more phased heterozygous sites genome-wide, with consistent improvements across long-read sequencing platforms (Figures 2, S1, & S2).

To our knowledge, LongHap is the first phasing method to selectively apply belief propagation to embed INDELs, SVs, and low-support variants into a broader haplotype context. This targeted embedding – combined with local realignment around complex breakpoints – improves phasing accuracy at challenging loci without incurring the computational overhead associated with globally embedding all variants, as reflected by LongHap’s ability to phase a large fraction of INDELs and SVs, while maintaining a low switch error rate and fast runtimes (Figures 2D, S1D, S2D). LongHap phases chromosome 1 of HG002 using 38x PacBio Revio HiFi data with and without methylation information in under ten minutes, substantially faster than MethPhaser, while remaining competitive with existing tools, with runtimes and memory scaling with sequencing depth and read length (Figures S6-S8).

By integrating methylation signals directly into the phasing process, LongHap extracts substantially more information than MethPhaser, which leverages methylation signals *post hoc* to refine pre-inferred haplotypes. While MethPhaser slightly increases switch error rates in our benchmarks, LongHap consistently reduces switch errors and extends phase blocks, demonstrating that joint modeling of genetic and epigenetic signals is more effective than *post hoc* refinement (Figures 2, S1, and S2).

Importantly, the benefits of methylation-aware phasing extend beyond genome-wide benchmarks to challenging, medically relevant genes and rare variants. Integrating methylation information reduces the number of phase blocks overlapping CMRGs by 11.3% and enables complete phasing of more CMRGs than any other tool, directly improving the interpretability of clinically important genomic regions (Table 1). For rare variants, which are systematically underpowered in statistical phasing approaches^8^, methylation integration consistently phases hundreds more rare variants per genome across individuals of diverse ancestries, with the largest gains in individuals of European and East Asian ancestry, where lower heterozygosity creates more frequent sequence-based phasing gaps. Critically, even modest improvement can have outsized clinical consequences since correctly resolving whether two rare variants occur in *cis* or *trans* determines their combined functional impact and is essential for interpreting compound heterozygosity in recessive disorders^5,17,18^. As exemplified by our analyses of two rare variants at the STXBP2 locus, LongHap can facilitate such analyses, aiding the interpretation of genetic variation in clinical settings.

By jointly leveraging information from read sequences and methylation patterns, LongHap provides a flexible framework that can be naturally extended to additional epigenetic or read-level signals, such as 5-hydroxymethylcytosine (5hmC), N6-methyladenine (6mA), and other base modifications that become detectable with routine long-read sequencing protocols in the future^33^. While orthogonal experimental assays such as chromatin conformation capture can provide long-range phasing information, they are not always practical in population-scale studies due to cost, sample availability, or experimental complexity. As long-read sequencing becomes increasingly prevalent, methods that maximize the information content of these data by integrating complementary signals will be essential.

While we showed that harnessing methylation information improves read-based variant phasing in individuals of diverse genetic ancestries, our analyses are based on sequencing data derived from an Epstein–Barr virus–transformed lymphoblastoid cell lines, which may exhibit atypical methylation patterns. However, the consistency of improvements observed across individuals of diverse ancestries suggests that haplotype-specific methylation patterns informative for phasing are a general feature of human genomes rather than a cell line-specific artifact. Nonetheless, additional studies of primary tissues will be useful to characterize the generalizability of methylation-aware phasing.

In summary, by jointly leveraging sequence and methylation information embedded in long-read sequencing data within a unified, probabilistic, and computationally efficient framework, LongHap substantially improves haplotype inference across variant classes and genomic contexts, including complex SVs, INDELs, and rare variants in challenging, medically relevant genes. These advances establish methylation-aware phasing as a broadly applicable strategy for studies of genetic variation, disease mechanisms, haplotype-specific epigenetic regulation, and human evolutionary history.

## Online Methods

### LongHap

#### Input for LongHap

As a minimal input, LongHap requires aligned sequencing reads in BAM format and variant calls in VCF format. To incorporate epigenetic information from long-read sequencing data, LongHap can optionally use site-specific methylation calls, generated, for example, with pb-CpG-tools and provided as unphased methylation annotations.

#### Determining allelic support in individual reads

Using pysam v0.22.1 (https://github.com/pysam-developers/pysam) and cyvcf2 v0.31.1^34^, LongHap streams through the BAM file and infers the allele each read supports at individual overlapping heterozygous sites in the VCF. Alignments with a mapping quality less than a specified threshold (by default, 20) are ignored along with secondary alignments, duplicate reads, and reads that are flagged as quality control failures. Unlike other read-based phasing tools, LongHap can include “multiallelic” heterozygous sites, where both haplotypes are different from the reference. On an individual genome level, these sites are still biallelic and simply require a dynamic indexing for accurate phasing. However, for comparability purposes, all described results are based on biallelic sites only.

To optimize runtime, LongHap first attempts to infer allelic states based on CIGAR strings and matching subsequences, resorting to local realignments only if the allele supported by a read cannot be confidently identified. For single-nucleotide variants (SNVs), LongHap requires an exact match of bases when using highly accurate PacBio Revio reads, and for noisier ONT reads, a matching base upstream and downstream of the heterozygous SNV is required in addition to a supporting CIGAR string. For insertions of any size, LongHap requires that the CIGAR string supports either no insertion/deletion (INDEL) or structural variant (SV) at all, or an insertion of the exact length as expected based on the VCF record, along with a matching inserted sequence. Similarly, for deletions, the CIGAR string must not support an INDEL, an SV of any size, or a deletion with the exact length specified in the VCF record. If the allelic state of a read at a given heterozygous site cannot be unambiguously identified according to the above criteria, the read-site pair is left for local realignment.

If a read-site pair is left for realignment, the read is locally re-aligned to synthetic references representing allele A and B, respectively. Synthetic references are created by embedding both alleles in the surrounding reference sequence. For PacBio, SNVs, INDELs, and SVs are embedded in 33 bps, 100 bps, and 100 bps, respectively. For noisier ONT data, SNVs, INDELs, and SVs are embedded in 66 bps, 200 bps, and 200 bps, respectively. However, the embedding size can also be defined from the command line for specific use cases. Gap open and extension penalties are dynamically set based on sequencing technology, variant type, and sequence context. Using the CIGAR string, we calculate the INDEL rate for each read. That is, we count the number of bases implicated in an insertion or deletion operation and divide it by the total number of CIGAR operations. For PacBio HiFi sequencing, the indel rate is typically less than 0.005, and we set the gap open and extension penalty to seven and two, respectively. For ONT sequencing, the indel rate can be higher depending on the kit used. For reads with indel rates, *r*, satisfying 0.005 ≤ *r* < 0.02, we set the gap open and extension penalties to 5 and 1, respectively. For reads with 0.02 < *r* ≤ 0.05, we set the gap open penalty to 4, and for reads with *r* > 0.05, we set the gap open penalty to 3, while keeping the gap extension penalty set to 1. Independent of the sequencing technology, we then double the gap open penalty for SNVs. Similarly, because both PacBio and ONT sequencing have higher error rates in homopolymeric sequences, we reduce the gap open penalty by 2 if a heterozygous site falls within a homopolymer, which we define as at least two identical bases surrounding the heterozygous site in both directions. Finally, we realign the read against both synthetic reference sequences, using a semiglobal alignment approach implemented in parasail v1.3.4 (*sg_stats_striped_sat*)^35^. We then assume that a read supports the allele yielding the greater alignment score.

#### Construction of directed-acyclic graph

After determining the allelic states of reads at overlapping heterozygous sites, we construct a directed acyclic graph (DAG) 𝒢 = (*V, E, ω*). *V*, the set of vertices, is defined by the heterozygous sites in the VCF file, including INDELs and SVs. For each heterozygous site, we create a layer with two vertices, representing two alleles (*A* and *B*), such that:

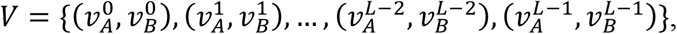

where *L* is the number of heterozygous sites in the VCF. *E*, the set of edges, comprises all possible edges between vertices of adjacent layers in 𝒢, that is:

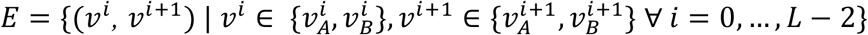

where 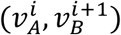 represents the edge connecting allele *A* and *B* at site *i* and *i* + 1, respectively. Edge weights, *ω*, are calculated by tallying the co-occurrence of alleles at two adjacent heterozygous sites *i* and *i* + 1, calculating the frequency with which allele pairs (*A*_*i*_, *A*_*i*+1_), (*A*_*i*_, *B*_*i*+1_) etc., co-occur at sites *i* and *i* + 1. The weights of edges connecting layers *i* and *i* + 1 (*ω*(*v*^*i*^, *v*^*i*+1^)) are thus defined by:

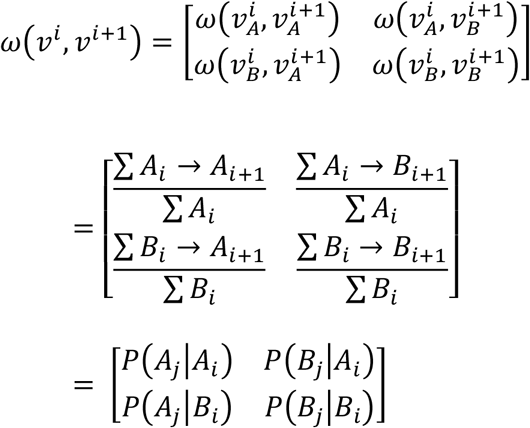

Finally, LongHap mirrors the more confident transition, creating a symmetrical 2*x*2 matrix for each pair of adjacent heterozygous sites.

When analyzing ONT data, we require that each allele is observed on both DNA strands since it was observed that variants supported by reads from only one DNA strand are more frequently false-positive calls^13^. Variants that are only observed on reads from one strand are marked as not phaseable and are excluded from subsequent analyses.

#### Re-estimating transitions for difficult-to-phase variants using loopy belief propagation

To improve haplotype reconstruction of difficult-to-phase variants, that is, INDELs, SVs, and variants for which at least one allele was observed less than *AN* times (*AN* < 1 by default), LongHap embeds them into the broader haplotype context by considering long-range edges between non-adjacent heterozygous sites. Note that alleles observed zero times (*AN* = 0) can infrequently occur as the result of alignment and variant calling errors or difficulty in determining the allelic state of individual reads by LongHap. To re-phase *m* consecutive difficult-to-phase variants (*i*, …, *i* + *m*), a subgraph 𝒢_𝒮_ = (*V*_*S*_, *E*_*S*_, *ω*_*S*_) is constructed with the vertices *V*_*S*_ = {*i* − *n*, …, *i*, …, *i* + *m*, …, *i* + *m* + *n*}, where *n* is the number of surrounding variants that can be confidently phased, and *E* = {(*v*^*j*^, *v*^*j*+1^)|*j* ∈ *i* − *n*, … *i* + *m* + *n*}. In practice, we choose *n* = 2. Each vertex can take one of two states *x* = {*A, B*}. For each pair of adjacent vertices, we already have a 2*x*2 transition matrix *ω*(*v*^*i*^, *v*^*i*+1^), indicating probabilities with that vertex *i* transitions from its current state into either state at vertex *i* + 1. In addition to these edges between adjacent pairs, long-range edges are considered, connecting the preceding and succeeding confidently phased vertices with all other vertices in the graph that are more than one step away, that is, *ω*(*v*^*j*^, *v*^*k*^) | *j* ∈ {*i* − *n*, …, *i* − 1} ∀ *k* ∈ {*j* + 2, …, *j* + *m* + *n* + 1} and *ω*(*v*^*j*^, *v*^*k*^) | *j* ∈ {*i*, …, *i* + *m* − 1} ∀ *k* ∈ {*j* + 2, …, *j* + *m* + *n* − 1}. For example, if *m = 1* and *n=2*, long-range transitions are inferred between vertex *i* − 2 and vertices *i* … *i* + 2, vertex *i* − 1 and vertices *i* + 1 and *i* + 2, and vertex *i* and vertex *i* + 2.

Belief Propagation (BP) provides an efficient means to infer marginal distributions of states at individual vertices by factorizing the graph and propagating previously calculated messages between vertices^36,37^. In acyclic graphs, BP yields an exact solution, while in graphs with cycles, only an approximation can be obtained. Although the above-described directed subgraph has no cycles *a priori*, it does have cycles in its moralized (undirected) form. We thus must approximate marginal distributions at individual vertices, using loopy BP^37,38^.

Given the above-described moralized subgraph, LongHap determines the neighbors for each vertex based on whether they are connected by an edge. Furthermore, states for each vertex *j* (*ϕ*_*j*_) are initialized uniformly, except for the first vertex (*i* − *n*), which is initialized to be in state *A*. The message a node *j* sends to a neighbor *j* (*m*_*j*→*k*_ (*x*_*k*_)) depends on its state *ϕ*_*j*_, the messages it receives from its other neighbors, and the local factor connecting *j* and *k* (*ψ*_*j,k*_(*x*_*j*_, *x*_*k*_) = *ω*(*v*^*j*^, *v*^*k*^)):

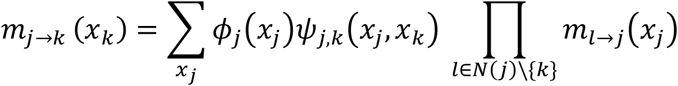

The loopy BP is started at an arbitrary pair of vertices *j* and *k*, and an updated 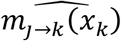 calculated. Based on 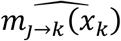, all other messages in the graph are updated, and the algorithm is run iteratively until it converges or reaches a maximum number of iterations (by default, 500). Although convergence is not guaranteed, we found that it virtually always converges within a few iterations. Given the messages received by a node *j*, its normalized marginal belief (*b*_*j*_(*x*_*j*_)) is given by:

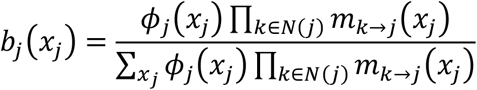

Similarly, unnormalized pairwise marginal beliefs of vertices *j* and *k* are given by:

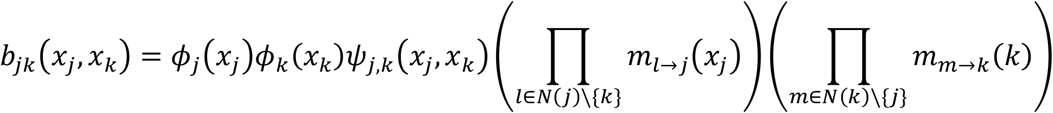

*b*_*jk*_ (*x*_*j*_, *x*_*k*_) can be interpreted as the transition probability between *j* and *k*. However, we are only interested in 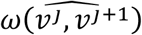 of adjacent heterozygous sites in the graph that is stripped of the long-range connections but implicitly still contains their information, that is, the information of long-range connections is projected in pairwise adjacent connections. 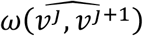 is thus obtained by solving:

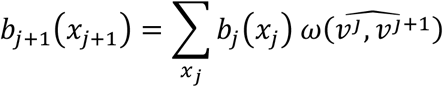

yielding:

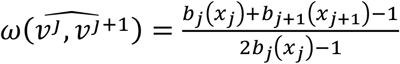

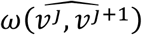 is consistent with the full DAG described above. In the example graph shown in Figure 1B, LongHap aims to calculate 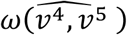 and 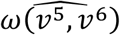 as described above.

#### Initial attempt to resolve ambiguous transitions

Although long-read sequencing allows phasing of most pairs of adjacent heterozygous sites, not all transitions can be unambiguously resolved. For some pairs *i* and *i* + 1, all transitions between allelic states may have equal probability, causing a break in the phase block (see below). In such cases, LongHap attempts to extend the phase block by testing whether the transitions from *i* to *i* + 2 can be confidently inferred. If this is the case, LongHap tests if it can use this long-range edge to update the posterior beliefs for the transitions from *i* to *i* + 1 and *i* to *i* + 2, or marks the heterozygous variant at site *i* + 1 as not phaseable and connects *i* to *i* + 2.

#### Leveraging methylation information to resolve remaining ambiguous transitions

After constructing the initial DAG based on sequence evidence, LongHap leverages methylation information inherent to PacBio and ONT long-read sequencing data to resolve remaining ambiguous transitions. These include pairs of adjacent heterozygous sites *i* and *i* + 1 that are not overlapped by any read, making it impossible to phase them solely based on sequence evidence.

To leverage methylation calls, LongHap first identifies putatively differentially methylated sites, that is, sites at which 20-80% of the reads are methylated. A haplotype is considered methylated if the product of methylation probabilities across all reads assigned to the haplotype is greater than 0.5, and its log10 odds ratio of being methylated versus unmethylated is greater than three, and unmethylated otherwise. Finally, a site is considered differentially methylated and informative for phasing if the inferred methylation states differ between the two haplotypes. LongHap uses these differentially methylated sites as additional markers to connect to phase blacks.

LongHap anchors reads based on their allelic states at sites *i* − 1 and *i* and then tests if parental haplotypes are differentially methylated at any candidate sites overlapped by these reads as described above. It then uses the differentially methylated sites to iteratively assign reads to either haplotype and identify additional differentially methylated sites based on the newly assigned reads. This procedure is repeated until either the gap between sites *i* and *i* + 1 is bridged or no more reads can be assigned to either haplotype. If the gap was successfully bridged, LongHap uses all reads assigned to a haplotype to infer the transition probabilities between sites *i* and *i* + 1 (*ω*(*v*^*i*^, *v*^*i*+1^)).

#### Viterbi-like inference of most likely haplotype

After completing the DAG, LongHap employs a Viterbi-like decoding scheme to identify the most likely haplotypes. That is, there are just transition probabilities and no emission probabilities in our model. At the beginning of the forward pass, we assign equal probabilities of being in state *A* and *B* at the first layer. LongHap then calculates the probability of being in either state at each subsequent layer of the DAG, keeping track of the more likely state. Whenever it encounters an ambiguous transition between two layers, that is, all transitions are equally likely, it stops the forward pass, randomly initializes a new phase block, and backtraces the previous phase block.

#### Output

LongHap writes a phased VCF, providing a PS tag for each phased variant. Optionally, LongHap also provides phase block coordinates, haplotagged read alignments in BAM format (HP tag), and information on the methylation sites used for phasing.

### Sequencing data

We retrieved multiple PacBio and ONT sequencing datasets for HG002 for benchmarking purposes from the T2T consortium^19^. Specifically, we downloaded PacBio Revio HiFi reads, ONT Simplex R10.4.1 reads base-called with Dorado (PAW70337), and UL-ONT Duplex R10.4.1 reads base-called with Dorado. We also retrieved PacBio Revio HiFi sequencing data for six additional samples from HPRC: two East Asian samples (HG00609, HG00658), two admixed samples from the Americas (HG00738, HG01099), and two sub-Saharan African samples (HG02723, HG02615)^20^. Table S1 summarizes the characteristics of the sequencing datasets used and provides links to each dataset.

### Sequence alignment, variant calling, and methylation calling

Sequences were aligned to CHM13v2.0 using minimap2 v2.28-r1221-dirty with --*map-hifi* and *--map-ont* presets for PacBio Revio HiFi and ONT sequencing data, respectively^39,40^. We then used PacBio Revio HiFi sequence alignments to call SNVs and small INDELs with DeepVariant v1.9.0 and larger SVs with Sniffles2 v2.7.1^41,42^. We removed sites with a genotype quality less than 20 (GQ < 20) and merged DeepVariant and Sniffles2 variant calls. Finally, we left-aligned and merged multiallelic variants using *bcftools norm -m+* ^43^. We determined methylation states using *aligned_bam_to_cpg_scores* v3.0.0 from pb-CpG-tools (https://github.com/PacificBiosciences/pb-CpG-tools).

### Benchmarking of read-based variant phasing methods

We compared the performance of LongHap to WhatsHap v2.3^9,11^, HapCUT2 v1.3.4^10^, LongPhase^13^, and WhatsHap + MethPhaser v0.0.3^16^. For each tool, we calculated phasing statistics using *whatshap stats* and switch error rates relative to the T2T HG002 Q100 variant call set using *whatshap compare*^9,11,19^. Prior to evaluation, variant calls in the T2T HG002 Q100 ground truth call set and phased variant calls were left-aligned, and multiallelic variants were split using *bcftools norm -m -any*^43^. As these tools phased variable percentages of heterozygous sites, we calculated switch error rates based on sites phased by all tools to allow a fair comparison.

We also evaluated the ability of each tool to contiguously phase a recently curated list of 273 challenging, medically relevant genes (CMRGs) made available by the Genome in a Bottle consortium by intersecting their gene bodies with phase blocks using bedtools^21,44^. Although some tools support parallelization, we used single CPU threads for all cores to enable fair comparisons of computational requirements.

In addition to HG002, we compared LongHap’s variant phasing for six additional individuals from Africa, East Asia, and the Americas to the phasing in the respective assemblies from HPRC release 2 (Table S1)^20^, using WhatsHap as described above. To obtain a ground truth set, we called variants for each assembly with dipcall v0.3^45^. We then annotated variant calls for each individual with allele frequencies from the 1000 Genomes Project to specifically evaluated LongHap’s ability to phase rare variants (MAF < 1%)^40^.

## Data availability

No new data has been generated. All used data is publicly available from the website of the T2T consortium and the Human Reference Pangenome Consortium and is summarized in Table S1.

## Code availability

LongHap and all code for supplementary analyses are available from: https://github.com/AkeyLab/LongHap.

## Acknowledgements

We thank Rob Biermann for providing computational support, helping to improve the computational footprint of LongHap, and all members of the Akey lab for insightful discussions. We also acknowledge that the work reported on in this paper was substantially performed using high-performance computing resources at the Lewis-Sigler Institute for Integrative Genomics and the Princeton Research Computing resources at Princeton University. Princeton Research Computing is a consortium of groups, including the Princeton Institute for Computational Science and Engineering (PICSciE) and Research Computing at Princeton University.

## Supplemental information

Figures S1–S8 and Tables S1 and S3

