## Supplemental Materials for "Methylation-aware long-read phasing significantly improves genome-wide haplotype reconstruction"

**The PDF file includes:**

Supplementary Figures S1-S8
Supplementary Tables S1-S3

### Supplementary Figures

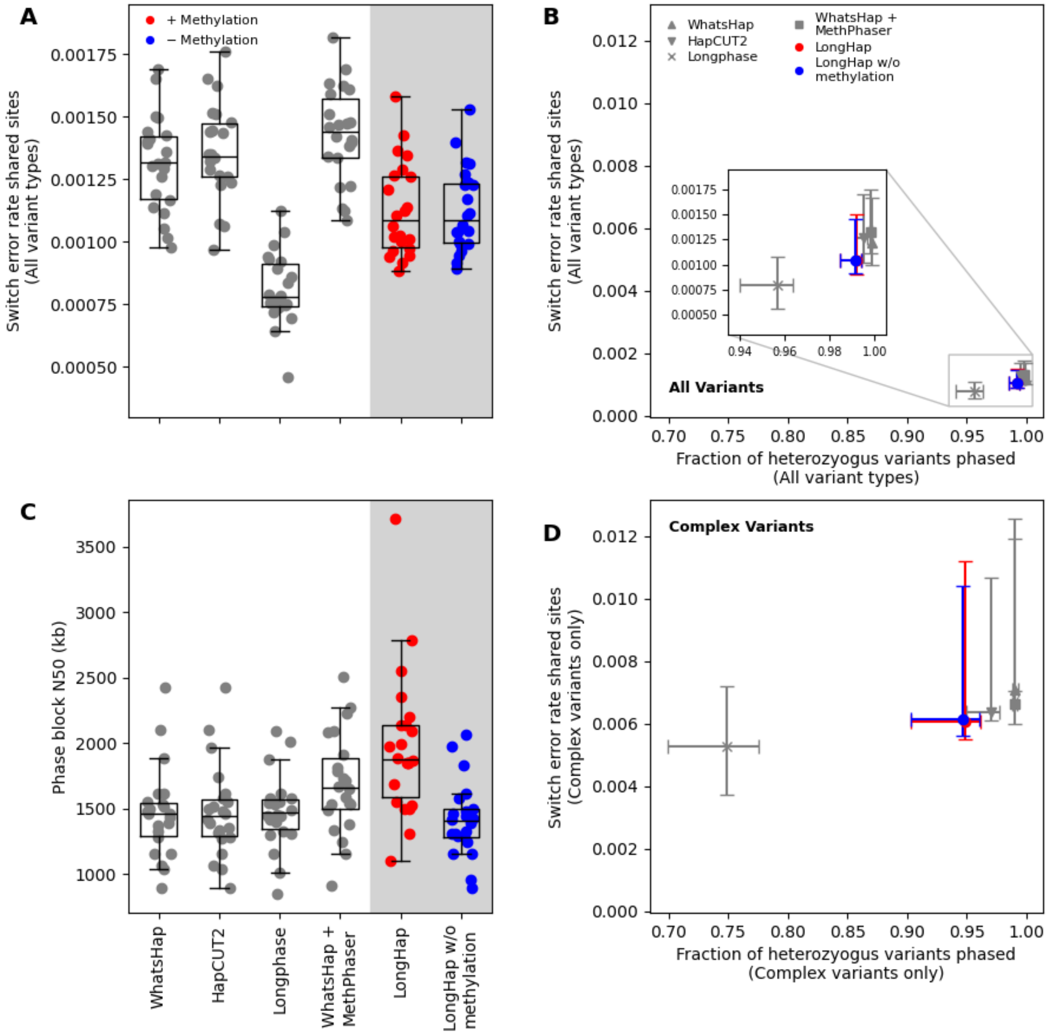


**Figure S1. Leveraging methylation information reduces switch error rates, including for INDELs and SVs, and increases phasing contiguity. A)** Switch error rates for different read-based variant phasing tools using 45x ONT R10.4.1 sequencing data base-called with Dorado for individual chromosomes of the well-characterized genome of HG002. LongHap is sensitive to the greater noise in ONT sequencing data but still achieves lower switch error rates than WhatsHap, HapCUT2, and WhatsHap + MethPhaser. Switch error rates were calculated based on the intersection of sites phased by all (93.77% of all heterozygous sites). **B)** LongHap phases, on average, 99.17% and 99.20% of all sites without and with methylation information – more sites than LongPhase (on average, 95.66%) – while maintaining a low error rate, through the integration of methylation information and its rigorous embedding of INDELs and SVs into the broader haplotype context. Switch error rates were calculated based on the intersection of sites phased by all tools (93.77% of all heterozygous sites with variants). **C)** Simultaneously, LongHap's integration of methylation information significantly increases the phase block N50 (mean phase block N50s: 1.97 Mb compared to 1.42 Mb without methylation information). **D)** As **(B)** but for INDELs and SVs. LongHap phases more INDELs and SVs than LongPhase, while maintaining a low error rate, through the integration of methylation information and its rigorous embedding of INDELs and SVs into the broader haplotype context. Switch error rates were calculated based on the intersection of INDEL and SV sites phased by all tools (70.17% of all heterozygous sites with complex variants).

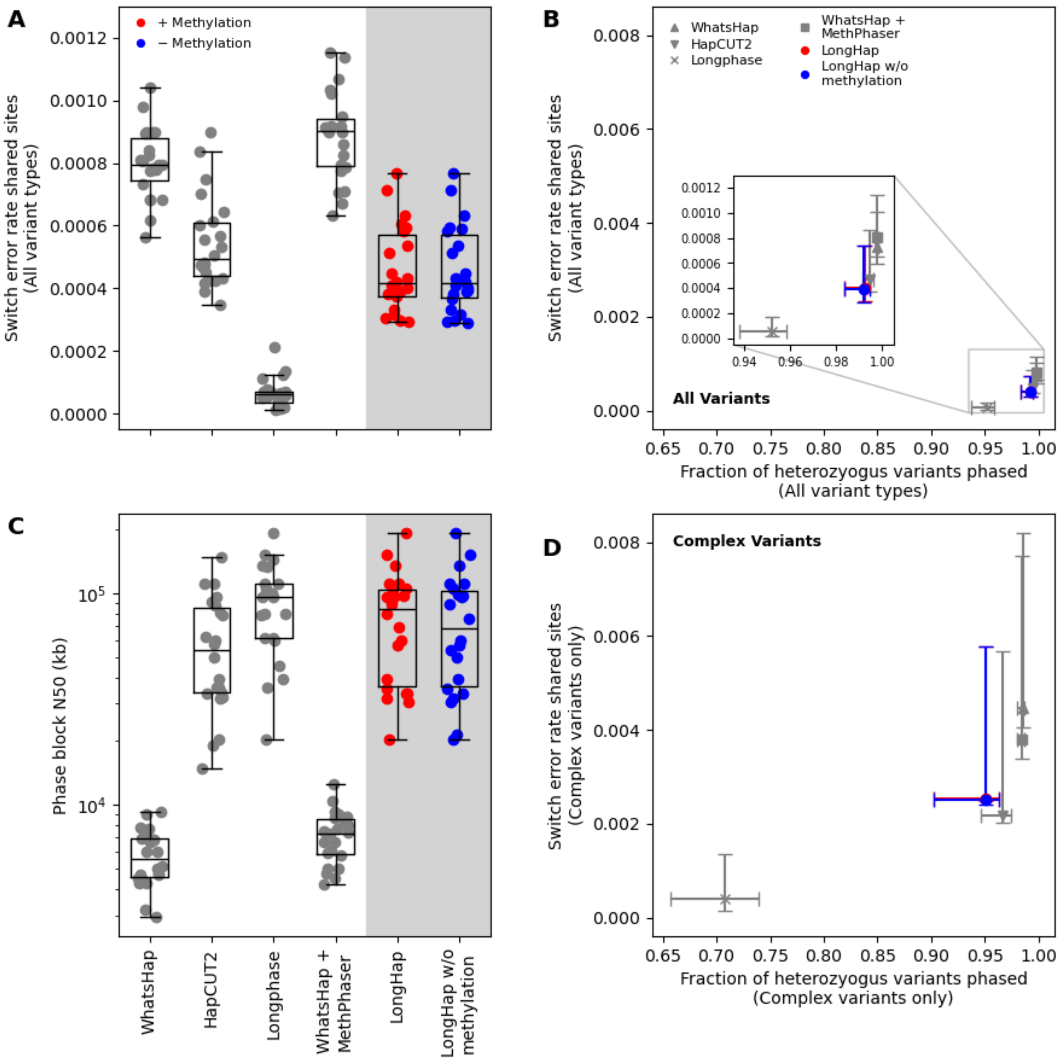


**Figure S2. Leveraging methylation information reduces switch error rates, including for INDELs and SVs, and increases phasing contiguity. A)** Switch error rates for different read-based variant phasing tools using 44x UL-ONT R10.4.1 sequencing data base-called with Dorado for individual chromosomes of the well-characterized genome of HG002. LongHap is sensitive to the greater noise in ONT sequencing data but still achieves lower switch error rates than WhatsHap, HapCUT2, and WhatsHap + MethPhaser. Switch error rates were calculated based on the intersection of sites phased by all (93.33% of all heterozygous sites). **B)** LongHap phases, on average, 99.22% of all sites without and with methylation information – more sites than LongPhase (on average, 95.21%) – while maintaining a low error rate, through the integration of methylation information and its rigorous embedding of INDELs and SVs into the broader haplotype context. Switch error rates were calculated based on the intersection of sites phased by all tools (93.33% of all heterozygous sites with variants). **C)** Simultaneously, LongHap's integration of methylation information increases the phase block N50, achieving chromosome-scale phasing (mean phase block N50s: 80.7 Mb compared to 77.1 Mb without methylation information). Note that the y-axis is shown in log scale. **D)** As **(B)** but for complex variants, that is,INDELs and SVs. LongHap phases more INDELs and SVs than LongPhase, while maintaining a low error rate, through the integration of methylation information and its rigorous embedding of INDELs and SVs into the broader haplotype context. Switch error rates were calculated based on the intersection of INDEL and SV sites phased by all tools (66.37% of all heterozygous sites with complex variants).

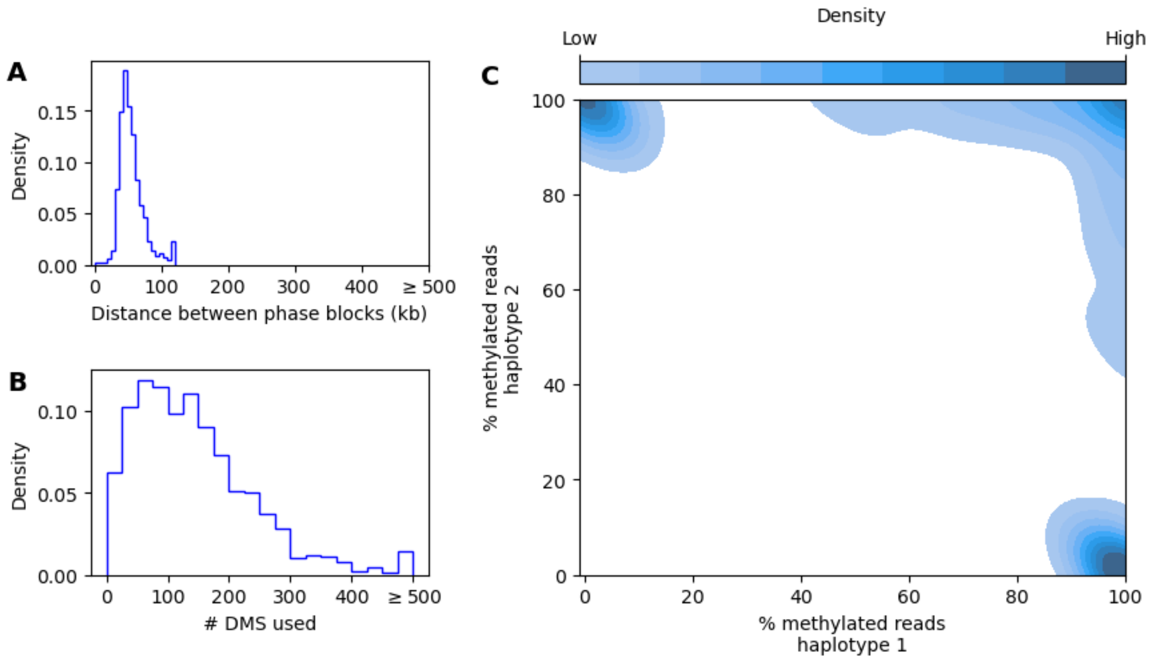


**Figure S3. Characterization of differentially methylated sites (DMS) used by LongHap for gap-bridging. A)** Distribution of distances between phase blocks connected using methylation information, using 45x ONT R10.4.1 sequencing data base-called with Dorado. Most bridged gaps are between 30–90 kb, with a small number of gaps exceeding 120 kb. **B)** Distribution of the number of DMS sites used to bridge each gap. Most gaps are resolved using approximately 100 DMS sites, with diminishing frequency at higher counts. **C)** Contour plot the fraction of reads with methylation marks assigned to either haplotype at sites used for phasing by LongHap. Most sites show clear signals of differential methylation, that is, they are truly phase-informative, as most reads are methylated on one haplotype while most reads are not methylated on the other haplotype (top left and bottom right corners). However, LongHap also identifies some false-positive differentially methylated sites (top right corner), injecting noise into the methylation-based phasing. A read is methylated at a given site if $P(meth) > 0.5$.

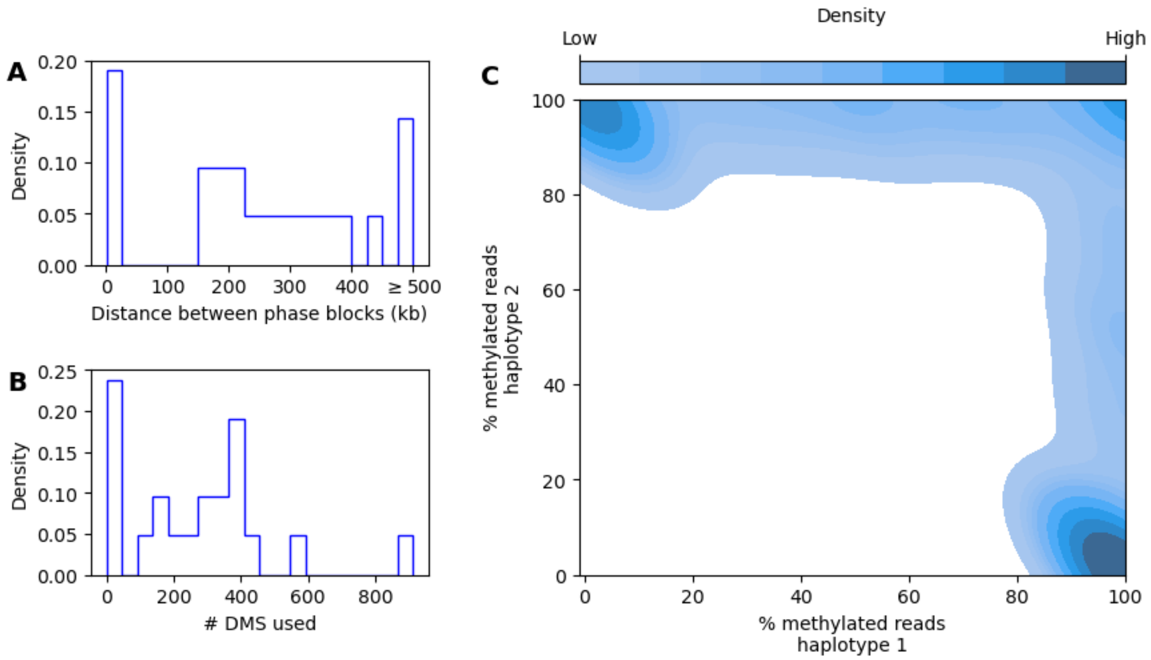


**Figure S4. Characterization of differentially methylated sites (DMS) used by LongHap for gap-bridging. A)** Distribution of distances between phase blocks connected using methylation information, using 44x UL-ONT R10.4.1 sequencing data base-called with Dorado. Most bridged gaps are between 150–400 kb, but ~15% also exceed 500 kb. **B)** Distribution of the number of DMS sites used to bridge each gap. Most gaps are resolved using approximately 400 DMS sites, with diminishing frequency at higher counts. **C)** Contour plot the fraction of reads with methylation marks assigned to either haplotype at sites used for phasing by LongHap. Most sites show clear signals of differential methylation, that is, they are truly phase-informative, as most reads are methylated on one haplotype while most reads are not methylated on the other haplotype (top left and bottom right corners). However, LongHap also identifies some false-positive differentially methylated sites (top right corner), injecting noise into the methylation-based phasing. A read is methylated at a given site if $P(meth) > 0.5$.

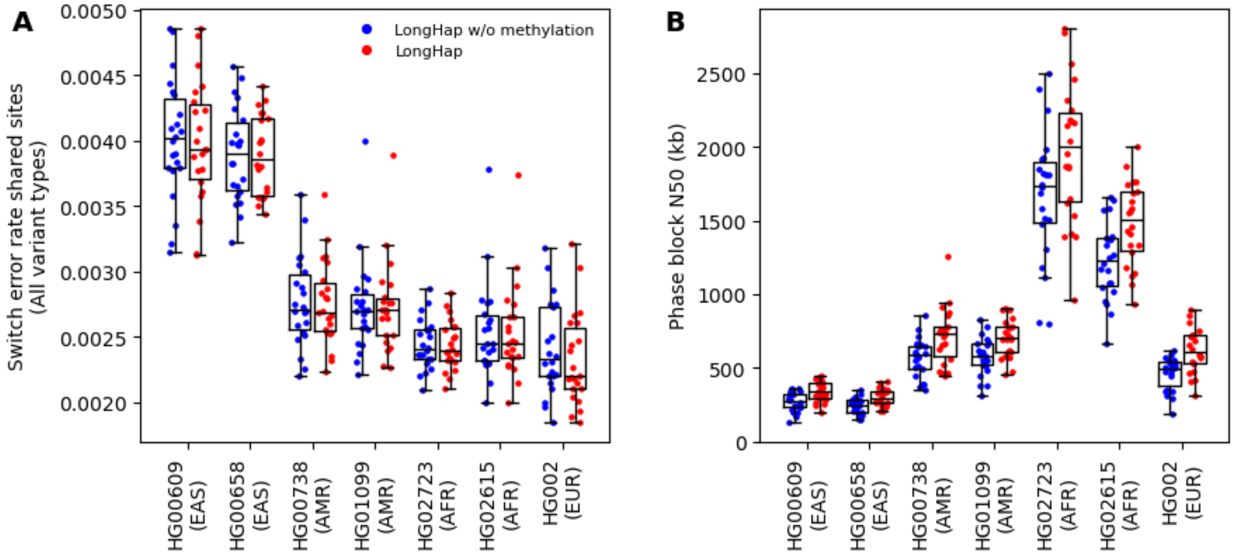


**Figure S5. Leveraging methylation information improves variant phasing across genetic ancestries. A)** Switch error rates for LongHap with and without methylation information using PacBio Revio HiFi sequencing data for individual chromosomes of seven individuals from diverse genetic ancestries (East Asian (EAS), admixed American (AMR), African (AFR), and European (EUR)). Switch error rates were calculated relative to the respective assemblies from release 2 of the Human Pangenome Reference Consortium. **B)** Phase block N50s for LongHap without (blue) and with methylation (red) information. While LongHap's performance differs between individuals depending on levels of heterozygosity, integrating methylation information improves switch errors and phase block contiguity by similar relative amounts, independent of genetic ancestry.

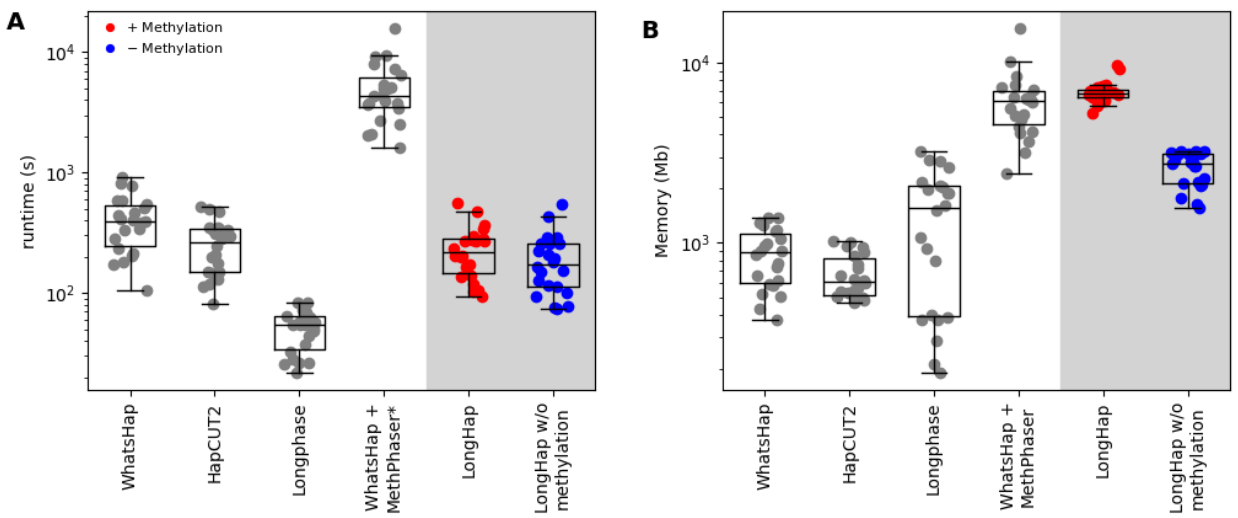


**Figure S6. LongHap integrates methylation information with little computational overhead using 38x PacBio Revio HiFi data. A)** Runtime in seconds and **B)** memory requirements in Mb for each tool for individual chromosomes of the well-characterized genome of HG002. LongHap requires between 74 and 547 seconds per chromosome when not integrating methylation information and between 94 and 557 seconds with methylation information. This is faster than WhatsHap (106 – 912 seconds), HapCUT2 (82 – 522 seconds), and MethPhaser (1,624 – 15,810 seconds without counting the runtime for WhatsHap and additional preparation of input files), but slower than LongPhase (22 – 84 seconds). LongHap requires modest amounts of memory, ranging between 1,562 Mb and 3,234 Mb without methylation information and 5241 Mb and 9,710 Mb when integrating methylation information. Note that all y-axes are shown in log scale.

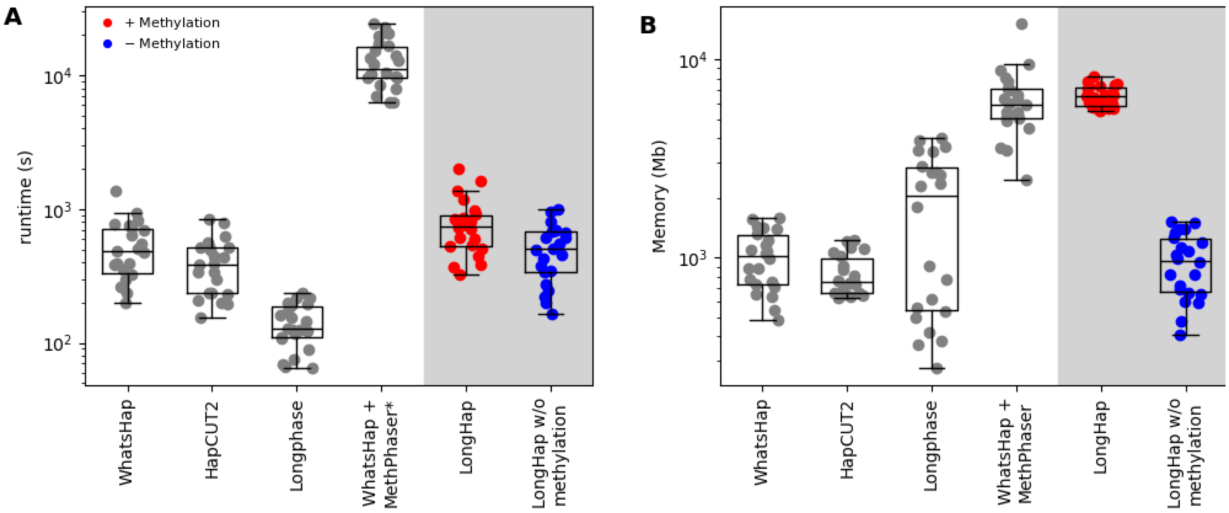


**Figure S7. LongHap integrates methylation information with little computational overhead using 45x ONT R10.4.1 sequencing data base-called with Dorado. A)** Runtime in seconds and **B)** memory requirements in Mb for each tool for individual chromosomes of the well-characterized genome of HG002. LongHap requires between 164 and 992 seconds per chromosome when not integrating methylation information and between 325 and 2,017 seconds with methylation information. This is similar to WhatsHap (165 – 1,140 seconds) and HapCUT2 (145 – 748 seconds), faster than MethPhaser (6,226 – 24,395 seconds without counting the runtime for WhatsHap and additional preparation of input files), but slower than LongPhase (62 – 748 seconds). LongHap requires modest amounts of memory, ranging between 408 Mb and 1,513 Mb without methylation information and 5,478 Mb and 8,143 Mb when integrating methylation information. Note that all y-axes are shown in log scale.

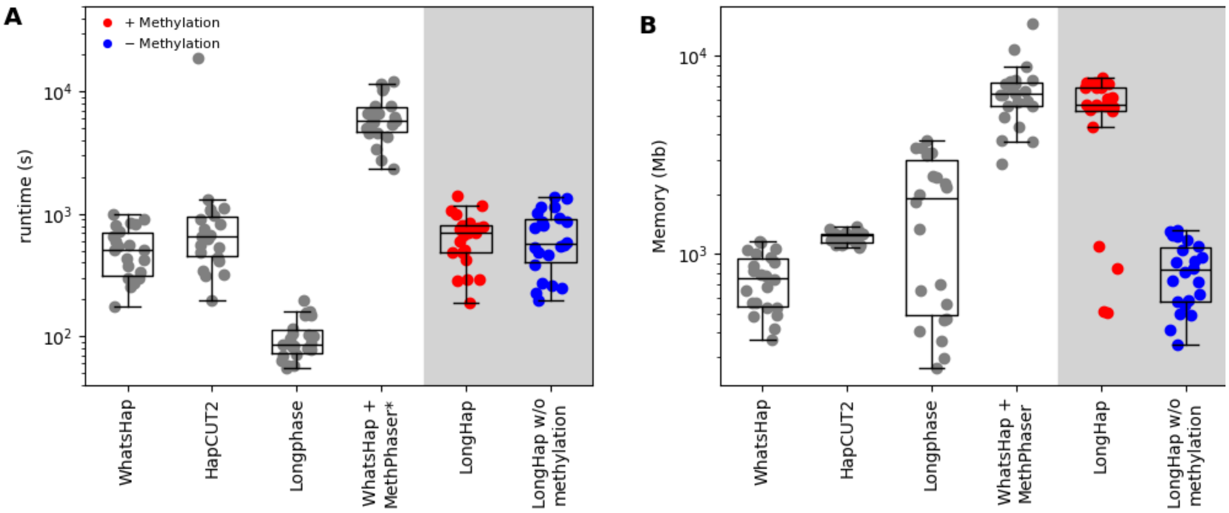


**Figure S8. LongHap integrates methylation information with little computational overhead using 44x UL-ONT R10.4.1 sequencing data base-called with Dorado. A)** Runtime in seconds and **B)** memory requirements in Mb for each tool for individual chromosomes of the well-characterized genome of HG002. LongHap requires between 197 and 1,365 seconds per chromosome when not integrating methylation information and between 191 and 1,410 seconds with methylation information. This is similar to WhatsHap (233 – 1,448 seconds) and HapCUT2 (199 – 18,551 seconds), faster than MethPhaser (2,351 – 12,126 seconds without counting the runtime for WhatsHap and additional preparation of input files), but slower than LongPhase (44 – 192 seconds). LongHap requires modest amounts of memory, ranging between 347 Mb and 1,314 Mb without methylation information and 501 Mb and 7,729 Mb when integrating methylation information. Note that all y-axes are shown in log scale.

#### Supplementary Tables

**Table S1:** Sequencing datasets used in this study and their characteristics.

| Sample | Sequencing dataset | Coverage | Read length N50 (kb) | Base quality (Q) | URL |
|---|---|---|---|---|---|
| **HG002** | **PacBio Revio HiFi** | ~38x | 18 | >30 | https://human-pangenomics.s3.amazonaws.com/submissions/80d00e88-7a92-46d8-88c7-48f1486e11ed--HG002_PACBIO_REVIO/m84039_230117_233243_s1.hifi_reads.default.bam |
| **HG002** | **ONT Simplex R10.4.1, Dorado** | ~45x | 29 | ~26 | s3://ont-open-data/giab_2025.01/basecalling/sup/HG002/PAW70337/calls.sorted.bam |
| **HG002** | **UL-ONT Duplex R10.4.1, Dorado** | ~44x | 111 | N/A | https://s3-us-west-2.amazonaws.com/human-pangenomics/index.html?prefix=T2T/scratch/HG002/sequencing/ont/11_16_22_R1041_UL_HG002_dorado0.4.0/ |
| **HG00609** | **PacBio Revio HiFi** | ~30x | 19 kb | >30 | s3://human-pangenomics/working/HPRC/HG00609/raw_data/PacBio_HiFi/m84081_231124_014115_s1.hifi_reads.bc2090.bam, s3://human-pangenomics/working/HPRC/HG00609/raw_data/PacBio_HiFi/m84081_231124_021221_s2.hifi_reads.bc2090.bam, s3://human-pangenomics/working/HPRC/HG00609/raw_data/PacBio_HiFi/m84081_231124_024327_s3.hifi_reads.bc2090.bam |
| **HG00658** | **PacBio Revio HiFi** | ~28x | 18 kb | >30 | s3://human-pangenomics/working/HPRC/HG00658/raw_data/PacBio_HiFi/m84091_230905_192642_s1.hifi_reads.bc1018.bam, s3://human-pangenomics/working/HPRC/HG00658/raw_data/PacBio_HiFi/m84091_230905_195701_s2.hifi_reads.bc1018.bam, s3://human-pangenomics/working/HPRC/HG00658/raw_data/PacBio_HiFi/m84091_230905_202807_s3.hifi_reads.bc1018.bam |
| **HG00738** | **PacBio Revio HiFi** | ~30x | 20 kb | >30 | s3://human-pangenomics/working/HPRC/HG00738/raw_data/PacBio_HiFi/m84081_231112_034048_s4.hifi_reads.bc2079.bam |

| | | | | | |
|---|---|---|---|---|---|
| **HG01099** | **PacBio Revio HiFi** | ~26x | 21 kb | >30 | s3://human-pangenomics/working/HPRC/HG01099/raw_data/PacBio_HiFi/m84081_231124_014115_s1.hifi_reads.bc2092.bam, s3://human-pangenomics/working/HPRC/HG01099/raw_data/PacBio_HiFi/m84081_231124_021221_s2.hifi_reads.bc2092.bam, s3://human-pangenomics/working/HPRC/HG01099/raw_data/PacBio_HiFi/m84081_231124_024327_s3.hifi_reads.bc2092.bam |
| **HG02723** | **PacBio Revio HiFi** | ~36x | 19 kb | >30 | s3://human-pangenomics/submissions/548dd68a-3e67-44a6-8b39-954b7a8eb835--HPRC_REVIO_EA_2023/bc2012-HG02723/m84036_230317_175945_s2.hifi_reads.bc2012.bam, s3://human-pangenomics/submissions/548dd68a-3e67-44a6-8b39-954b7a8eb835--HPRC_REVIO_EA_2023/bc2012-HG02723/m84039_230303_012244_s3.hifi_reads.bc2012.bam, s3://human-pangenomics/submissions/548dd68a-3e67-44a6-8b39-954b7a8eb835--HPRC_REVIO_EA_2023/bc2012-HG02723/m84039_230314_213047_s2.hifi_reads.bc2012.bam, s3://human-pangenomics/submissions/548dd68a-3e67-44a6-8b39-954b7a8eb835--HPRC_REVIO_EA_2023/bc2012-HG02723/m84039_230316_193003_s2.hifi_reads.bc2012.bam |
| **HG02615** | **PacBio Revio HiFi** | ~36x | 18 kb | >30 | s3://human-pangenomics/working/HPRC/HG02615/raw_data/PacBio_HiFi/m84081_231105_031800_s4.hifi_reads.default.bam |

**Table S2.** Number of phase blocks inferred by each tool using 45x ONT R10.4.1 sequencing data base-called with Dorado that overlap challenging, medically relevant genes (CMRGs). 243 CMRGs are covered by all tools. LongHap infers the most contiguous phase blocks overlapping CMRGs, as indicated by the lowest number of phase blocks and the second most overlapping base pairs.

| | CMRGs covered | Phase blocks overlapping CMRGs | CMRGs fully phased | Phase blocks overlapping CMRGs covered by all tools | Overlapping Base pairs of CMRGs covered by all tools (Mb) |
|---|---|---|---|---|---|
| **LongHap** | 263 | 408 | 248 | 399 | 143.2 |
| **LongHap w/o methylation information** | 259 | 441 | 243 | 427 | 140.0 |
| **LongPhase** | 263 | 455 | 246 | 445 | 147.1 |
| **WhatsHap** | 262 | 482 | 245 | 469 | 139.1 |
| **HapCUT2** | 262 | 469 | 245 | 450 | 138.3 |
| **WhatsHap + MethPhaser** | 262 | 524 | 245 | 513 | 138.7 |

**Table S3.** Number of phase blocks inferred by each tool using 44x UL-ONT R10.4.1 sequencing data base-called with Dorado that overlap challenging, medically relevant genes (CMRGs). 256 CMRGs are covered by all tools. LongHap infers the most contiguous phase blocks overlapping CMRGs, as indicated by the lowest number of phase blocks and the second most overlapping base pairs.

| | CMRGs covered | Phase blocks overlapping CMRGs | CMRGs fully phased | Phase blocks overlapping CMRGs covered by all tools | Overlapping Base pairs of CMRGs covered by all tools (Mb) |
|---|---|---|---|---|---|
| **LongHap** | 273 | 278 | 271 | 256 | 157.3 |
| **LongHap w/o methylation information** | 272 | 278 | 270 | 256 | 157.1 |
| **LongPhase** | 272 | 273 | 271 | 256 | 175.7 |
| **WhatsHap** | 266 | 357 | 257 | 256 | 146.6 |
| **HapCUT2** | 271 | 284 | 268 | 256 | 154.2 |
| **WhatsHap + MethPhaser** | 266 | 387 | 256 | 256 | 146.1 |
